# Human spindle variability

**DOI:** 10.1101/2021.09.02.458632

**Authors:** Christopher Gonzalez, Xi Jiang, Jorge Gonzalez-Martinez, Eric Halgren

## Abstract

In humans, sleep spindles are 10-16 Hz oscillations lasting approximately 0.5-2 s. Spindles, along with cortical slow oscillations, facilitate memory consolidation by enabling synaptic plasticity. Early recordings of spindles at the scalp found anterior channels had overall slower frequency than central-posterior channels. This robust, topographical finding led to dichotomizing spindles as ‘slow’ versus ‘fast’, modelled as two distinct spindle generators in frontal versus posterior cortex. Using a large dataset of intracranial sEEG recordings (n=20, 365 bipolar recordings), we show that the difference in spindle frequency between frontal and parietal channels is comparable to the variability in spindle frequency within the course of individual spindles, across different spindles recorded by a given site, and across sites within a given region. Thus, fast and slow spindles only capture average differences that obscure a much larger underlying overlap in frequency. Furthermore, differences in mean frequency are only one of several ways that spindles differ. For example, compared to parietal, frontal spindles are smaller, tend to occur after parietal when both are engaged, and show a larger decrease in frequency within-spindles. However, frontal and parietal spindles are similar in being longer, less variable, and more widespread than occipital, temporal, and Rolandic spindles. These characteristics are accentuated in spindles which are highly phase-locked to posterior hippocampal spindles. We propose that rather than a strict parietal-fast/frontal-slow dichotomy, spindles differ continuously and quasi-independently in multiple dimensions, with variability due about equally to within-spindle, within-region and between-region factors.

**Significance Statement:** Sleep spindles are 10-16 Hz neural oscillations generated by cortico-thalamic circuits that promote memory consolidation. Spindles are often dichotomized into slow-anterior and fast-posterior categories for cognitive and clinical studies. Here, we show that the anterior-posterior difference in spindle frequency is comparable to that observed between different cycles of individual spindles, between spindles from a given site, or from different sites within a region. Further, we show that spindles vary on other dimensions such as duration, amplitude, spread, primacy and consistency, and that these multiple dimensions vary continuously and largely independently across cortical regions. These findings suggest that multiple continuous variables rather than a strict frequency dichotomy may be more useful biomarkers for memory consolidation or psychiatric disorders.

## Introduction

Brain rhythms during sleep play an active role in organizing and strengthening our memories (Marshall et al., 2020; Rasch and Born, 2013). Two such rhythms are cortico-thalamic slow waves and sleep spindles. Slow waves are large, ∼0.5-4Hz rhythms composed of alternating downstates (periods of neuronal quiescence), followed by upstates, where neuronal activity is similar to waking (Steriade et al., 1993). Spindles are 10-16 Hz oscillations lasting 0.5-2s, and are initiated by the inhibitory thalamic reticular nucleus interacting with excitatory thalamocortical cells (Steriade, 2003; Steriade et al., 1993). Spindles are grouped by slow waves (Mak-Mccully et al., 2017), facilitating memory consolidation (Mölle et al., 2009; Niknazar et al., 2015). Synaptic plasticity during spindles may be enabled by dendritic calcium influx (Seibt et al., 2017; Sejnowski and Destexhe, 2000), and neuronal co-firing (Dickey et al., 2021).

Early scalp recordings (Gibbs and Gibbs, 1950) distinguished slower spindles at frontal sensors from faster spindles at posterior sensors. This observation has promoted a model of spindle dynamics as two distinct spindle generators: slow-frontal and fast-parietal (Anderer et al., 2001; Ayoub et al., 2012; Mölle et al., 2011; Timofeev and Chauvette, 2013). Alternatively, many local spindle generators with overlapping frequency distributions could be spread across the cortex (Dehghani et al., 2011a, 2010; Frauscher et al., 2015; Gennaro and Ferrara, 2003; Peter-Derex et al., 2012; Piantoni et al., 2017). In this case, frequency differences between frontal and parietal sites would not reflect two generators with uniform frequency, but rather the average differences of generators that have highly overlapping frequencies in each location. In this study, we quantify the degree of frequency variation between regions, within a region, at individual cortical sites, and within individual spindles, to better adjudicate between these two generating mechanisms.

Frequency variability within spindles includes a spectral-spatial shift over the course of the spindle. In both MEG and EEG, power is maximal at higher frequencies (13-15 Hz) earlier in the spindle, especially at central sensors, and maximal at lower frequencies (10-12 Hz) later in the spindle, especially at frontal sites (Dehghani et al., 2011a; Zygierewicz et al., 1999). Here we show that differences in intra-spindle frequency variability are large compared to differences in frequency due to region. Previous scalp (O’ Reilly and Nielsen, 2014; Schönwald et al., 2011; Souza et al., 2016) and intracranial (Andrillon et al., 2011) studies have also reported a systematic slowing during spindles. We provide a more comprehensive study in spindle slowing across the cortex, as well as report how more widespread spindles slow more than local spindles.

These findings support updating a model of cortical spindles from two spindle generators to a model with many generators with varied and overlapping frequency characteristics. Spindles vary in many characteristics besides frequency, including duration, spread, amplitude, waveform, and relationship to other waves within and outside the cortex. Such differences, if they follow the frontal-slow/parietal-fast differences, would reinforce that dichotomy. Indeed, the prime evidence for a biological distinction between slow and fast spindles is the assertion that slow precede and fast follow the downstate (Klinzing et al., 2016; Mölle et al., 2011). However, our previous findings indicate that all spindles follow (Gonzalez et al., 2018; Mak-Mccully et al., 2017). Here, we systematically explore variations in these characteristics and their correlations with each other, and the location where they are recorded. We found that these different characteristics vary quasi-independently, clearly indicating that cortical spindle generators cannot be divided into two distinct types located in different lobes, but rather form a mosaic of generators varying in multiple characteristics, often with large overlap across anatomical regions. This reconceptualization may inform future efforts to relate spindles to memory consolidation and clinical disorders.

## Methods

### Patient selection

Patients with intractable, pharmaco-resistant epilepsy were implanted with stereoelectroencephalographic (sEEG) electrodes to determine seizure onset for subsequent resection for treatment. Patients were selected from an original group of 54, excluding patients that had pronounced diffuse slowing, widespread interictal discharges, or highly frequent seizures. Patients were further selected that each had at least one hippocampal contact in a hippocampus not involved in seizure initiation. The remaining 20 patients include 7 males, aged 29.8 +/-11.9 years old (range 16-38 years). For demographic and clinical information, see Table 1 originally published in Jiang et al 2019a. All electrode implants and duration of recordings were selected for clinical purposes (Gonzalez-Martinez et al., 2013). All patients gave fully informed consent for data usage as monitored by the local Institutional Review Board, in accordance with clinical guidelines and regulations at Cleveland Clinic.

**Table 1.** List of patients, age, sex, handedness, NC channel counts, and the lengths of sleep period recordings used. Pt: patient. L: Left. R: Right. Dur.: duration. Std dev: standard deviation. T: Total. Reused with permission from Jiang et. al. 2019a.

| Pt # | Age | Sex | Hand-<br>edness | #<br>Cortical<br>Channels | HC<br>Seizure-<br>free? | HC<br>Interictal-<br>free? | # Sleep<br>Periods | mean<br>duration<br>of NREM<br>(h) | Std<br>dev<br>NREM<br>(h) | Total<br>N2<br>Dur.<br>(h) | Total<br>N3<br>Dur.<br>(h) |
| --- | --- | --- | --- | --- | --- | --- | --- | --- | --- | --- | --- |
| 1 | 20 | M | R | 19 | Y | N | 3 | 2.2 | 0.78 | 2.8 | 3.3 |
| 2 | 51 | F | R | 12 | Y | N | 4 | 7.5 | 1.87 | 27.3 | 0 |
| 3 | 58 | F | R | 24 | Y | N | 4 | 7.1 | 2.44 | 23.1 | 0.30 |
| 4 | 42 | M | L | 17 | Y | N | 4 | 3.1 | 0.33 | 1.8 | 9.0 |
| 5 | 18 | F | L | 21 | Y | Y | 1 | 3.7 |  | 1.9 | 0.9 |
| 6 | 20 | F | R | 21 | Y | N | 3 | 2.7 | 1.49 | 2.6 | 3.0 |
| 7 | 22 | M | LR | 19 | Y | Y | 3 | 3.8 | 1.17 | 2.4 | 3.8 |
| 8 | 30 | F | R | 13 | Y | N | 5 | 5.2 | 1.26 | 6.5 | 14 |
| 9 | 43 | F | R | 12 | Y | N | 4 | 3.1 | 0.45 | 3.7 | 4.4 |
| 10 | 16 | M | R | 17 | Y | Y | 5 | 3.8 | 1.11 | 7.5 | 8.8 |
| 11 | 32 | F | R | 30 | Y | N | 3 | 5.1 | 3.43 | 8.4 | 2.8 |
| 12 | 36 | M | L | 25 | Y | Y | 4 | 5.2 | 1.48 | 11.6 | 6.5 |
| 13 | 21 | F | L | 14 | Y | N | 3 | 6 | 0.47 | 14.7 | 1.1 |
| 14 | 21 | F | R | 14 | Y | N | 8 | 3.7 | 1.1 | 16.9 | 9.3 |
| 15 | 29 | F | R | 17 | Y | N | 4 | 2.5 | 1.31 | 5.4 | 2.7 |
| 16 | 41 | F | R | 19 | Y | N | 3 | 4.6 | 0.72 | 7.4 | 4.4 |
| 17 | 24 | M | R | 21 | Y | N | 3 | 5.1 | 1.88 | 8.9 | 2.9 |
| 18 | 31 | F | R | 15 | Y | N | 6 | 6.1 | 1.18 | 24.2 | 3.8 |
| 19 | 21 | M | R | 15 | Y | Y | 4 | 2.8 | 0.72 | 5.4 | 5.8 |
| 20 | 19 | F | R | 21 | Y | Y | 3 | 4.9 | 2.53 | 5.2 | 6.0 |
| mean | 30 |  |  | 18 | T:20 | T:8 | 4 | 4.4 |  | 9.39 | 4.64 |

### Electrode localization

Electrodes were localized by registering a post-operative CT scan with a pre-operative 3D T1-weighted MRI with ∼1mm^3^ voxel size (Dykstra et al., 2012) using Slicer (RRID:SCR_005619). Recordings were obtained for 32 HC contacts, 20 anterior (11 left) and 12 posterior (7 left). The CT-visible cortical contacts were identified as previously described (Jiang et al., 2019a), to ensure activity recorded by bipolar transcortical pairs is locally generated (Mak-McCully et al., 2015). Electrode contacts were excluded if they were involved in early stages of seizure discharge or had frequent interictal activity. Of the 2844 contacts implanted in the selected 20 patients, 366 transcortical pairs (18.3 +/-4.7 per patient) were accepted for further analysis. Polarity was adjusted, if necessary, to “pial surface minus white matter” according to MRI localization, confirmed with decreased high gamma power for surface-negative downstates.

Freesurfer was used to reconstruct pial and white matter surfaces from individual MRI scans (Fischl, 2012) and to parcellate the cortical surface into anatomical areas (Desikan et al., 2006). An average surface from all 20 patients was generated to serve as the basis for all 3D maps. Each transcortical contact pair was assigned anatomical parcels from the Desikan atlas by ascertaining the parcel identities of the surface vertex closest to the contact-pair midpoint. Transcortical contact pair positions for all patients were registered to the fsaverage template brain for visualization by spherical morphing (Fischl et al., 1999). Because some contact pair midpoints are buried within sulci, for visualization, all markers were moved to the same plane along one dimension for medial and lateral surfaces separately using MATLAB 2018a. This allows visualizing all contact markers while maintaining anatomical fidelity. For a priori statistical analysis of spindle characteristics by cortical region, insular transcortical pairs were assigned to temporal cortex, and paracentral, postcentral and precentral labels constituted the Rolandic cortex.

### Data processing

Continuous recordings from SEEG depth electrodes were made with a cable telemetry system (JE-120 amplifier with 128 or 256 channels, 0.016-3000 Hz bandpass, Neurofax EEG-1200,Nihon Kohden) across multiple nights (Table 1). Patients were recorded over the course of clinical monitoring for spontaneous seizures, with 1000Hz sampling rate. The total NREM sleep durations vary across patients due to intrinsic variability and sleep deprivation due to clinical environment. We confirmed the percentages of NREM in total sleep from 28 sleeps across 16 of our patients were comparable to (i.e. within 2 SD of) normative data (Moraes et al., 2014) in terms of stage 2 (N2) and stage 3 (N3) durations. Furthermore, we did not observe any significant differences in sleep graphoelements (GE) compared with normative data (Jiang et al., 2019a). Recordings were anonymized and converted into the European Data Format. Subsequent data processing was performed in MATLAB 2018a (RRID:SCR_001622); the Fieldtrip toolbox (Oostenveld et al., 2011) was used for line noise removal and visual inspection. The separation of patient NREM sleep/wake states from intracranial LFP alone was achieved by previously described methods using clustering of first principal components of delta-to-spindle and delta-to-gamma power ratios across multiple LFP-derived signal vectors (Gervasoni, 2004; Jiang et al., 2017), with the addition that separation of N2 and N3 was empirically determined by the proportion of downstates that are also part of slow oscillations (at least 50% for N3) (Silber et al., 2007), since isolated downstates in the form of K-complexes are predominantly in N2 (Cash et al., 2009). The data was collected during a selected period where patients were off anti-epileptic drugs, minimizing the effects of medication in the generation and morphology of sleep rhythms.

### Cortical graphoelement detection

Spindles were detected as previously reported (Gonzalez et al., 2018; Jiang et al., 2019a; Mak-McCully et al., 2017). For each sleep period, each channel’s signal was filtered in 4-8 Hz, 10-16 Hz, and 18-30 Hz bands using a zero-phase shift frequency domain filter. The width of the filter transition bands relative to the cut-off frequencies was 0.3 and a Hanning window was used for the transition. To calculate the power envelope for each narrow band signal, the absolute value of the filtered data was calculated and smoothed by convolution with a 400ms Tukey window. To detect peaks in the power time series, this signal was subsequently smoothed using a 600ms Tukey window and a robust estimate of deviation for each channel was calculated by subtracting the median and dividing by the median absolute deviation. Putative spindle peaks were identified as exceeding 3 in the normalized median power time series and a relative edge threshold of 40% of the peak amplitude defined spindle onsets and offsets. Spindle epochs that co-occurred with greater than 3 in theta (4-8 Hz) and beta (18-30 Hz) ranges were excluded.

Spindle detections were performed on each sleep period separately. Only spindles longer than 0.5s were analyzed.

Downstates and theta bursts were detected as previously described (Gonzalez et al., 2018; Jiang et al., 2019b). DSs were detected on each channel as follows: (1)apply a zero-phase shift, eighth order (after forward and reverse filtering) Butterworth IIR bandpass filter from 0.1 to 4 Hz; (2) select consecutive zero crossings within 0.25-3 s; and (3) calculate amplitude trough between zero-crossings and retain only the bottom 20% of troughs. Theta bursts were detected as follows: (1) apply a zero-phase shift, eighth order (after forward and reverse filtering) Butterworth IIR bandpass filter from 5 to 8 Hz (range selected to minimize overlap with delta and spindle content); (2) calculate the mean of the Hilbert envelope of this signal smoothed with a Gaussian kernel (300ms window; 40ms sigma); (3) detect events with a +/-3 SD threshold for the peak and identify the start and stop times with a +/-1 SD threshold; (4) only include events with a duration between 400ms and 1s; and (5) for each bandpass filtered peak in a putative burst, calculate the preceding trough-to-peak deflection and only take events that have at least 3 peaks exceeding 25% of the maximum deflection.

### Spindle characteristics

Several spindle characteristics were estimated for each spindle, including: duration, overall frequency, frequency change, spindle frequency variability, high gamma power, maximal spindle amplitude, and waveform shape measures. The duration of the spindle is defined as the onset and offset of the spindle as reported in *Cortical graphoelement detection*. We applied a zero-phase shift, eighth order (after forward and reverse filtering) Butterworth IIR bandpass filter at 10-16Hz and extracted the detected spindles. To be consistent with reported waveform shape measures, detected spindles were cropped such that troughs started and ended the spindle epoch. This defined a cycle’s period as trough-to-trough. For each cycle, the narrowband trough-to-peak amplitude as calculated. For a cycle to be included, it needed to exceed 30% of the maximum trough-to-peak amplitude within the spindle. The frequency of a cycle was calculated by dividing the sampling rate by the trough-to-trough period (in samples) and limiting the frequency precision to two decimal places. The frequency of each spindle was calculated as the number of cycles surviving the amplitude threshold divided by the trough-to-trough duration of the spindle. To assess frequency variability within a spindle, the difference of the fastest and slowest cycle in Hz and the standard deviation of cycle frequencies was calculated. Frequency change per spindle was estimated using least squares. For each spindle a design matrix, A, was constructed as an intercept column and a column indicating the time of each peak since the first peak (in seconds) in the narrow-band signal. The frequency at each cycle was coded as the dependent variable, b, and the coefficients for frequency intercept and change was estimated using the MATLAB ‘A\b’ operation. For each channel this yielded a distribution of frequency slopes and a one-sample t-test was used to assess significant slowing or speeding (in Hz/s). These measures were only calculated for spindles with at least 4 cycles that exceeded 30% of the maximum amplitude. High gamma (70-190 Hz) power was calculated by taking the square of the magnitude of the analytic signal. The median of this power vector over the duration of each spindle was calculated.

Measures of waveform shape were computed using custom MATLAB scripts as defined by previous literature (Cole and Voytek, 2019). Each channel was band-passed filtered in a broadband signal of 1-30 Hz via finite-impulse response filters (duration minimum of 3 cycles of 10 Hz). This signal was band-pass filtered in 10-16 Hz in order to identify times of zero-crossings. These times were mapped back to the broadband signal, and peaks (troughs) were identified as the maxima (minima) within these zero-crossings. Spindle epochs started and ended with troughs to set the number of cycles equal to the number of peaks. Because peaks are marked as maxima on the broadband signal, they could be marked during steep broadband rises or falls and not reflect true spindle cycles. To mitigate these effects, two rejection criteria were applied to each cycle within a spindle before estimating spindle metrics: 1) the cycle deflection (µV), defined as the average of the trough-to-peak and peak-to-trough amplitude, must exceed 30% of the largest cycle deflection within a spindle, and 2) the amplitude of the rising (falling) phase must exceed 15% of the amplitude of the falling (rising) phase. Applying an average amplitude threshold (1) removes smaller cycles, and (2) removes cycles that may have large amplitude but fall on steep rises or falls in the broadband signal. This step avoids analyzing cycles that would otherwise have rise-decay-symmetry measures artifactually close to 0 or 1. These steps mitigate the influence from overlapping large, slower rhythms on the spindle waveform shape features. For each cycle, the rise-decay symmetry was defined as the proportion of time spent in the rise phase (from trough-to-peak) out of the total trough-to-trough time. The peak-trough symmetry was defined as the duration of each peak (determined by zero-crossings) over the total duration of each peak and each trough. For a spindle to be included in waveform shape analysis, we required a density of 8 Hz, and to estimate mean and standard deviation within a spindle, at least 5 good cycles. The maximum spindle amplitude reported is the maximum trough-to-peak deflection in the broadband (1-30 Hz) signal across all cycles.

For all measures, only spindles exceeding a duration of 0.5 s were analyzed. The average of all spindle characteristics was calculated for each channel, and only channels with at least 50 spindles were analyzed. The NREM, N2, and N3 spindle density (spindles per min) was also calculated.

### Statistical analysis

These spindle characteristics were estimated for each spindle and subsequently averaged across spindles for each channel and for major cortical lobes. All statistical analyses were performed using R (R Core Team 2020).We applied linear mixed-effects models (LMEMs) to account for measuring multiple cortical channels within patients. This was calculated using the *lme4* package (Bates et al, 2015) at the channel level as ‘ lmer (Dependent Var ∼ 1 + Independent Fixed Effect + (1|Patient),data)’.For example, when comparing differences across cortical regions, cortical region was the fixed effect of interest, patients coded as random intercepts, and each observation was the average measure for a channel. When evaluating associations between spindle characteristics, each observation was a spindle and a nested random effect structure was applied as follows:’ lmer (Dependent Var ∼ 1 + Independent Fixed Effect + (1|Patient/Channel),data)’. When comparing the start versus end of spindle features (waveform shape measures, frequency), for each channel, paired-t-tests assessed significant differences across spindles. Descriptive tables and summary results of mixed effects models were created using R packages *qwraps2* (DeWitt 2021) and *sjPlot* (Lüdecke 2021), respectively. Boxplots were generated using *ggplot2* (Wickham 2016).

### Widespread spindles

The number of cortical channels spindling was calculated for each sample, and co-spindling events were defined as the non-zero onset and offset periods. For each spindle, we determined the maximum number of channels with minimum 100 ms overlap. This value was expressed both as the number of cortical channels co-spindling, as well as the proportion of channels co-spindling for each patient. We also recorded whether a spindle was the leading or initiating spindle in a co-spindle event, as well as its latency to start from the beginning of the event (expressed in ms).

### Assessment of hippocampal-cortical phase-locking value

The active process of sleep contributing to memory consolidation and structuring depends on communication transfer between the hippocampus and cortex (Buzsaki, 1996; Rasch and Born, 2013). Previous work in our lab identified a subset of parietal channels that have large phase-locking value (PLV > 0.4) in spindles, especially during N2, with posterior hippocampal spindles (Jiang et al., 2019b). In the current study, significant phase-locking between HC and NC spindles was determined as reported in Jiang et al 2019b. This analysis focused on coupling of NC channels to posterior HC during N2. Briefly, for each NC-HC pair, spindles detected in both structures that overlapped for at least 1 spindle cycle (here 160ms) had phase-locking values (PLVs) (Lachaux et al., 1999) calculated over 3s trials centered on all hippocampal spindle event starts in NREM. We also computed PLV for the same NC-HC channel pairs over the same number of trials centered on random times in NREM to create a baseline estimate; and for each non-overlapping 50 ms time bin, a two-sample t test was performed between the actual PLV and the baseline estimate, with the resulting p values undergoing FDR correction. A given channel pair would be considered significantly phase-locking if: (1) > 40 trials were used in the PLV computation; or (2) at least 3 consecutive time bins yield post-FDR p values <0.05. In this study, we characterized differences in spindle characteristics at sites with high PLV (>0.4) compared to all other cortical sites.

## Results

### Data characterization

We analyzed intracranial SEEG recordings from twenty patients with pharmaco-resistant epilepsy, and their full demographic information can be found in our previous work (Jiang et al. 2019a,b). Only cortical channels and sleep periods free of epileptic activity were selected for spindle analysis. Information pertinent to this study, including number of bipolar cortical recordings, number of sleep periods, and hours in N2 and N3 are shown in Table 1. In total we recorded from 366 cortical sites, and as shown in Figure 1, we have broad coverage across the cortical surface, including both hemispheres and medial and lateral surfaces. Of these 366 sites, 5 were excluded due to anatomical labels that were assigned “Medial wall” or “unknown” and 4 excluded because they did not have at least 50 spindles with durations greater than 500ms, resulting in 357 cortical sites analyzed.

**Figure 1.**
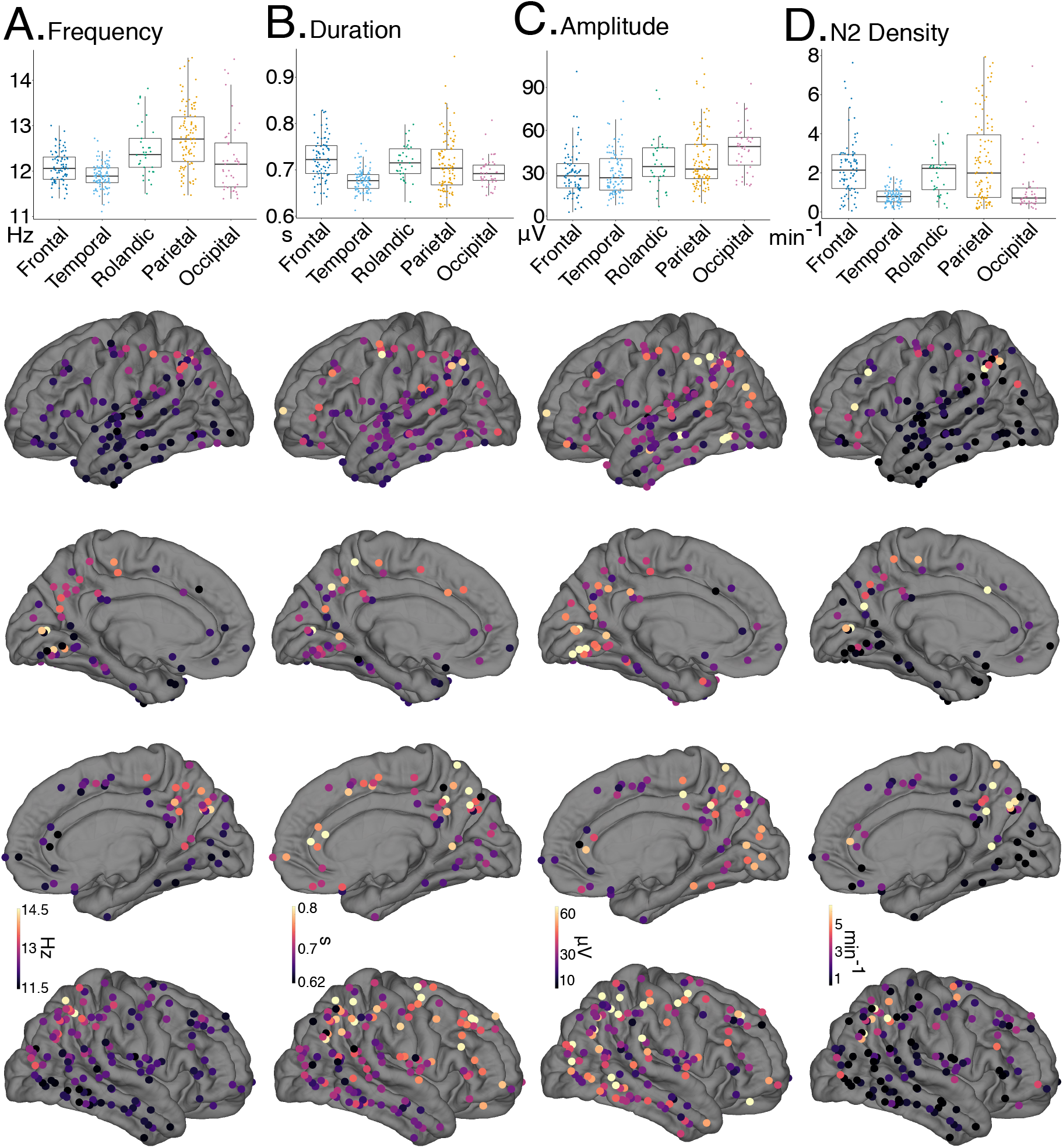
Primary spindle characteristics. Cortical bipolar SEEG recordings denoted as circles overlaid on an average surface, with warmer colors indicating greater values. Also shown at the top of the brain surfaces for each measure are boxplots grouped by brain region. For the boxplots, different colors and columns denote different regions, box margins indicate inter-quartile ranges, and dots indicate individual cortical channels. **A**. Average overall frequency at individual sites is shown. Frequency gradually increases from frontal-temporal to Rolandic to parietal. **B**. Average spindle duration across spindles at individual sites is shown. Compared to temporal and occipital, frontal-temporal-Rolandic structures have longer duration. **C**. Average maximal trough-to-peak voltage in a broadband (1-30Hz) signal increases along the anterior to posterior axis. **D**. Number of spindles per minute during stage 2 (N2), greatest for frontal and parietal sites.

### Primary spindle characteristics

Spindles were detected across the cortical surface in both N2 and N3 sleep. The average number of spindles and standard deviation across channels is reported in Table 2. The median number of spindles per channel and interquartile ranges for NREM, N2, and N3 are, respectively: 907 (357,2100), 649 (218,1315), and 209 (54,569). Spindle density, or number of spindles per minute, is shown for NREM sleep stages separately (Table 2). Regional differences in spindle density were observed, with frontal and parietal sites showing greater densities than temporal, Rolandic, or occipital areas (Figure 1D, Table2).

**Table 2.** Spindle characteristics for regions and non-REM sleep stage. Counts and means (SD) across channels shown.

|  | Cortex | Frontal | Temporal | Rolandic | Parietal | Occipital |
| --- | --- | --- | --- | --- | --- | --- |
| **Data** |  |  |  |  |  |  |
| No. Patients | 20 | 12 | 19 | 15 | 17 | 14 |
| No. Channels | 357 | 80 | 101 | 33 | 102 | 41 |
| **No. Spindles** |  |  |  |  |  |  |
| NREM | 1,531.90<br>(1,725.98) | 1,983.60<br>(1,342.65) | 717.16<br>(738.15) | 1,535.39<br>(1,018.27) | 2,015.47<br>(2,151.13) | 1,451.71<br>(2,527.84) |
| N2 | 1,035.79<br>(1,340.84) | 1,239.12<br>(949.93) | 582.27<br>(664.69) | 1,091.61<br>(823.01) | 1,280.22<br>(1,520.19) | 1,103.22<br>(2,444.47) |
| N3 | 496.11<br>(707.19) | 744.48<br>(658.88) | 134.89<br>(184.73) | 443.79<br>(494.25) | 735.25<br>(962.72) | 348.49<br>(591.08) |
| **Density (min<sup>-1</sup>)** |  |  |  |  |  |  |
| NREM | 1.64 (1.58) | 2.12 (1.45) | 0.71 (0.40) | 1.73 (1.10) | 2.30 (2.03) | 1.28 (1.60) |
| N2 | 1.82 (1.67) | 2.29 (1.50) | 0.85 (0.49) | 2.03 (1.20) | 2.55 (2.14) | 1.31 (1.56) |
| N3 | 1.43 (1.62) | 2.03 (1.53) | 0.49 (0.44) | 1.30 (1.11) | 2.08 (2.16) | 1.01 (1.13) |
| **Frequency (Hz)** |  |  |  |  |  |  |
| NREM | 12.28 (0.63) | 12.10 (0.34) | 11.89 (0.28) | 12.47 (0.59) | 12.74 (0.68) | 12.28 (0.80) |
| N2 | 12.30 (0.63) | 12.15 (0.36) | 11.90 (0.27) | 12.54 (0.59) | 12.75 (0.69) | 12.29 (0.79) |
| N3 | 12.22 (0.66) | 12.01 (0.34) | 11.84 (0.35) | 12.28 (0.63) | 12.73 (0.74) | 12.23 (0.77) |
| **Duration (s)** |  |  |  |  |  |  |
| NREM | 0.70 (0.05) | 0.72 (0.05) | 0.68 (0.02) | 0.72 (0.04) | 0.71 (0.06) | 0.69 (0.03) |
| N2 | 0.71 (0.05) | 0.73 (0.05) | 0.68 (0.03) | 0.72 (0.04) | 0.73 (0.07) | 0.70 (0.03) |
| N3 | 0.69 (0.06) | 0.72 (0.04) | 0.66 (0.03) | 0.72 (0.09) | 0.68 (0.06) | 0.69 (0.06) |
| **Amplitude (μV)** |  |  |  |  |  |  |
| NREM | 35.31<br>(18.19) | 30.14<br>(16.23) | 30.12<br>(14.69) | 36.90<br>(17.92) | 39.22<br>(20.24) | 47.19<br>(16.94) |
| N2 | 35.35<br>(18.16) | 30.31<br>(16.05) | 30.08<br>(14.58) | 36.73<br>(17.95) | 39.27<br>(20.27) | 47.16<br>(17.18) |
| N3 | 35.61<br>(18.41) | 30.47<br>(16.49) | 30.32<br>(14.71) | 38.69<br>(18.87) | 39.14<br>(20.52) | 47.18<br>(16.90) |

The average spindle amplitude and duration of spindles are shown in Table 2 as well as Figure 1. The average spindle amplitude (defined as maximum trough-to-peak deflection) increases anterior-to-posterior and is largest at occipital sites. Spindles have the longest duration at frontal, Rolandic, and parietal sites, and shortest at temporal sites.

The overall frequency of spindles showed a clear increase along the anterior-to-posterior axis, from frontal (12.1 Hz) to Rolandic (12.47 Hz) to parietal (12.74 Hz) regions, though showing intermediate frequencies at occipital sites (12.28 Hz). Temporal cortex showed the lowest overall frequency (11.89 Hz). Additionally, there appears to be a cluster of parietal channels with much higher frequency than surrounding cortex. These sites overlap with a subset of channels our group previously identified (Jiang et al 2019b) as showing strong phase-locking to hippocampal spindles and will be further discussed in section *Cortical-hippocampal spindle phase locking*.

### Variability in spindle frequency

In addition to spindle frequency varying between regions, we found there was substantial variability across sites within a region, across spindles measured at a single cortical site, and within individual spindles. Here we compare the magnitude of these sources of variability with our reported average difference between frontal and parietal channels of 0.64 Hz (12.10 vs. 12.74 Hz, Table 2), which is indicated with a black triangle on the scale bars in Figure 2.

**Figure 2.**
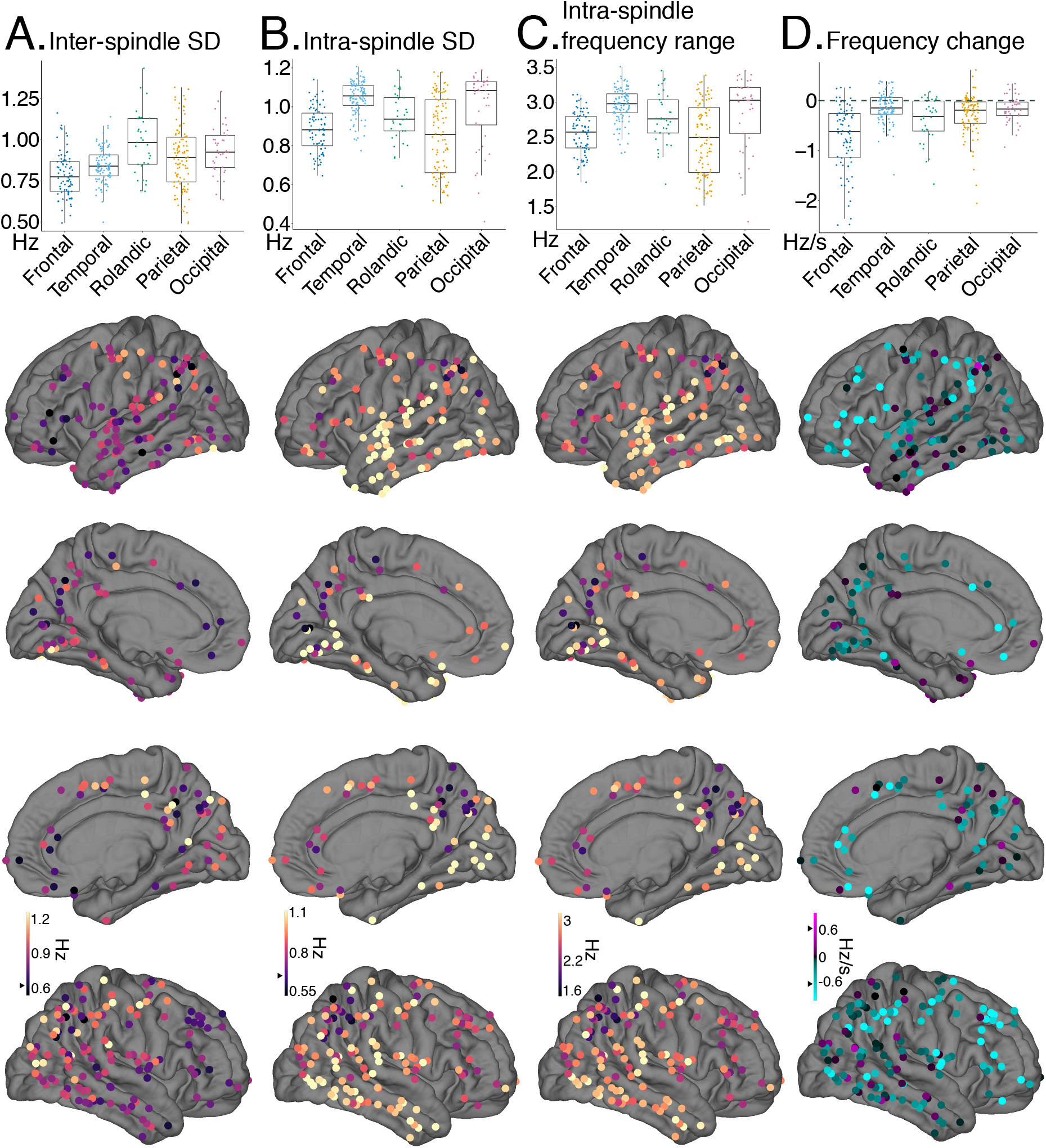
Sources of frequency variability. Cortical bipolar SEEG recordings denoted as circles overlaid on an average surface, with warmer colors indicating greater values. Black triangles mark 0.64 Hz (or Hz/s) to indicate the average difference between frontal and parietal sites. Also shown at the top of the brain surfaces for each measure are boxplots grouped by brain region. **A**. At individual cortical sites, the standard deviation of overall frequency across spindles. Rolandic sites show the greatest variability in frequency across spindles. **B**. The average standard deviation across cycle frequency within a spindle. Intra-spindle variability is largest for temporal and occipital sites. **C**. Another measure of intra-spindle frequency variation, the average difference between the fastest and slowest cycle within a spindle. **D**. Average estimated linear change in frequency within a spindle. The majority of cortical sites show a decrease in frequency or slowing during a spindle (cyan), with greatest slowing occurring at frontal sites. The amount of inter and intra-spindle variability often exceeds the observed frontal-parietal overall difference in frequency.

### Frequency variability across sites within a single cortical region

The variation within a region across channels was lower in frontal and temporal cortices, ranging from 0.28 to 0.34 Hz standard deviations respectively, compared to parietal and occipital sites ranging from 0.68 to 0.8 Hz respectively (see Fig1A boxplot). Notably, the standard deviation across channels within a site is larger at parietal and occipital channels than the average difference between frontal and parietal areas (0.64 Hz).

### Frequency variability at a single cortical site

To assess how much individual sites vary in spindle frequency, we calculated the standard deviation in overall frequency across spindles at each cortical channel (Fig 2A, Table 3A). Notably, the average inter-spindle frequency standard deviation is 0.87 Hz in NREM across the cortex. Rolandic sites had the largest inter-spindle standard deviation (0.98 Hz), whereas frontal sites had the lowest with 0.79 Hz. Thus the amount that spindles vary in frequency at a single site typically exceeds the reported difference between frontal and parietal recordings.

**Table 3.**
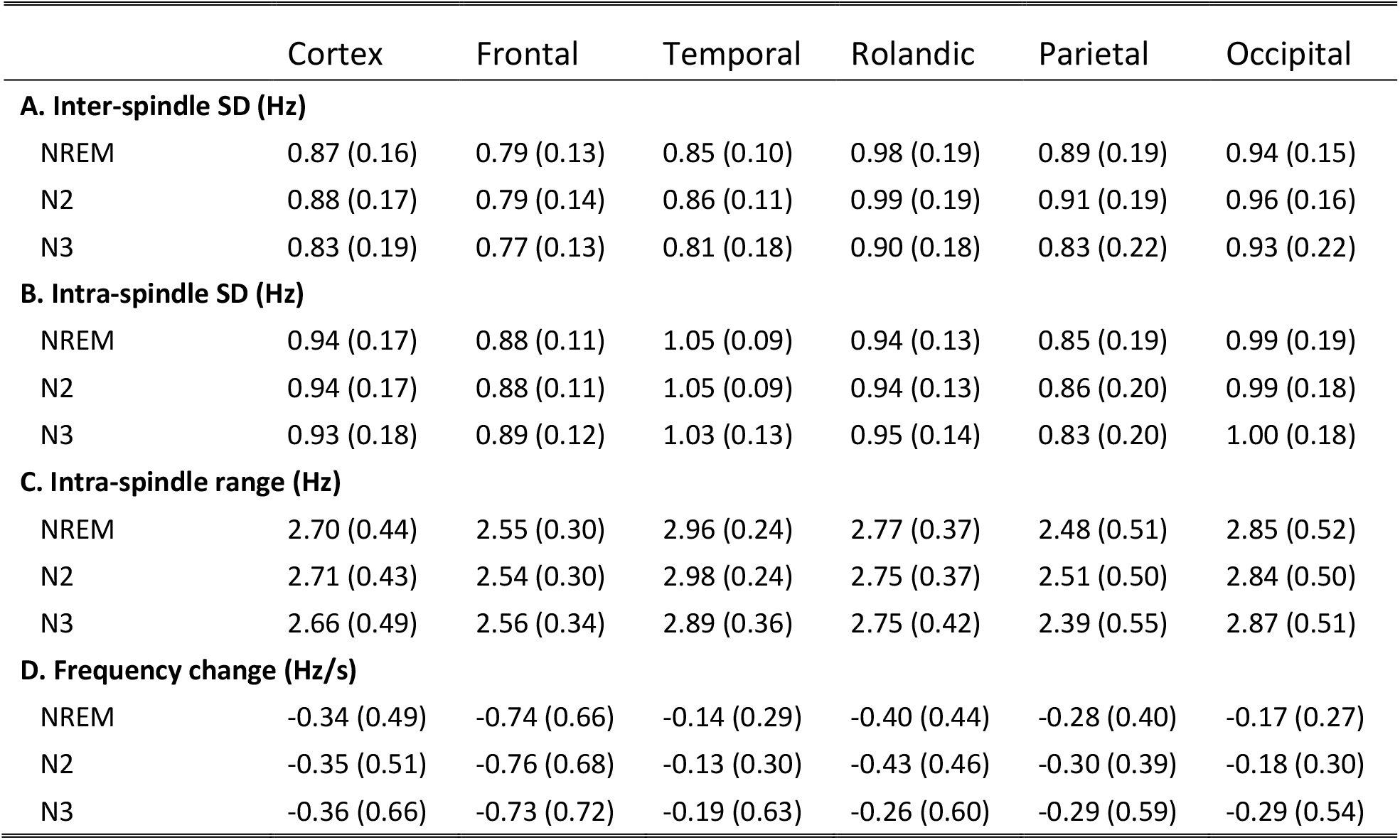
Spindle frequency variability. **A. Inter-spindle variability** as assessed by the mean SD across channels of the average frequency of its spindles, and the SD of this SD. **B. Intra-spindle variability** as assessed by first finding for each channel the mean SD of the cycle-by-cycle frequency across all of its spindles. Shown is the mean SD across all channels within an area, and the SD of this SD. **C. Intra-spindle range** as assessed by first finding for each channel the mean range of the cycle-by-cycle frequency across all of its spindles. Shown is the mean (SD) across channels within the area. **D. Frequency change** as assessed by first finding for each channel the mean slope of the cycle-by-cycle frequency across all of its spindles. Shown is the mean (SD) across channels within the area.

|  | Cortex | Frontal | Temporal | Rolandic | Parietal | Occipital |
| --- | --- | --- | --- | --- | --- | --- |
| **A. Inter-spindle SD (Hz)** |  |  |  |  |  |  |
| NREM | 0.87 (0.16) | 0.79 (0.13) | 0.85 (0.10) | 0.98 (0.19) | 0.89 (0.19) | 0.94 (0.15) |
| N2 | 0.88 (0.17) | 0.79 (0.14) | 0.86 (0.11) | 0.99 (0.19) | 0.91 (0.19) | 0.96 (0.16) |
| N3 | 0.83 (0.19) | 0.77 (0.13) | 0.81 (0.18) | 0.90 (0.18) | 0.83 (0.22) | 0.93 (0.22) |
| **B. Intra-spindle SD (Hz)** |  |  |  |  |  |  |
| NREM | 0.94 (0.17) | 0.88 (0.11) | 1.05 (0.09) | 0.94 (0.13) | 0.85 (0.19) | 0.99 (0.19) |
| N2 | 0.94 (0.17) | 0.88 (0.11) | 1.05 (0.09) | 0.94 (0.13) | 0.86 (0.20) | 0.99 (0.18) |
| N3 | 0.93 (0.18) | 0.89 (0.12) | 1.03 (0.13) | 0.95 (0.14) | 0.83 (0.20) | 1.00 (0.18) |
| **C. Intra-spindle range (Hz)** |  |  |  |  |  |  |
| NREM | 2.70 (0.44) | 2.55 (0.30) | 2.96 (0.24) | 2.77 (0.37) | 2.48 (0.51) | 2.85 (0.52) |
| N2 | 2.71 (0.43) | 2.54 (0.30) | 2.98 (0.24) | 2.75 (0.37) | 2.51 (0.50) | 2.84 (0.50) |
| N3 | 2.66 (0.49) | 2.56 (0.34) | 2.89 (0.36) | 2.75 (0.42) | 2.39 (0.55) | 2.87 (0.51) |
| **D. Frequency change (Hz/s)** |  |  |  |  |  |  |
| NREM | -0.34 (0.49) | -0.74 (0.66) | -0.14 (0.29) | -0.40 (0.44) | -0.28 (0.40) | -0.17 (0.27) |
| N2 | -0.35 (0.51) | -0.76 (0.68) | -0.13 (0.30) | -0.43 (0.46) | -0.30 (0.39) | -0.18 (0.30) |
| N3 | -0.36 (0.66) | -0.73 (0.72) | -0.19 (0.63) | -0.26 (0.60) | -0.29 (0.59) | -0.29 (0.54) |

### Frequency variability within a spindle

Next, we evaluated measures of intra-spindle variability. This includes the intra-spindle standard deviation, the range of spindle cycle frequency, and the linear change in frequency. The cortical average intra-spindle standard deviation, that is, across spindle cycles within a spindle, was 0.94Hz during NREM (Figure 2B; Table 3B). The average frequency range within a spindle, calculated as the difference of the fastest and slowest cycle, was 2.7 Hz (Figure 2C; Table 3C). For both of these measures, the temporal and occipital cortical sites had the largest intra-spindle variation, whereas the frontal and parietal had the lowest (Figure 2B,C; Table 3).

Intra-spindle variability is not completely random, but partially reflects a linear change in frequency. Previous work has reported that spindles decrease in frequency during their evolution. However, estimates across the cortex intracranially have not been systematically reported. 45.3% (163/360) of cortical channels showed spindle frequencies that were significantly different at the start versus at the end of the spindle (paired-t test, p<0.05, FDR adjusted). Of these, 98.2% (160/163) showed slower frequencies at the end versus the start. The three channels that showed faster frequencies at the end of the spindle were located in the lateral inferior temporal sulcus and pericalcarine cortex. For more precise estimates of linear change in frequency, we regressed cycle frequency against time since the first cycle for each spindle. The average change estimates across spindles for a given cortical site are shown in Figure 2D, and summarized in Table 3D. Using this approach, we found that 46.4% of cortical sites had a significant linear change in spindle frequency (one-sample t-test, p<0.05 FDR adjusted), with 95.2% (159/167) showing spindle slowing and 4.8% (8) channels showing spindle speeding. These 8 speeding channels were in the ventral and lateral temporal (2), pericalcarine (2), superior and inferior parietal (2), insula (1), and superior frontal cortex (1) across 7 different patients. As evident in Figure 2D, the majority of recordings across cortical regions tended to decrease in frequency, and significant slowing in spindle frequency occurred across frontal cortex (−0.74 Hz/s). To determine the proportion of overall intra-spindle frequency variation that is due to this consistent change in frequency, the observed frequency range within a spindle was compared with an estimated frequency range calculated by multiplying the average duration and frequency change per channel. On average, this estimated range accounted for 12% of the observed frequency range, with 75% of channels accounting for less than 14%. Thus, the average change in frequency across cycles of a given spindle is about three times larger than the average difference in frequency between frontal and parietal spindles, and only about 12% of the within-spindle variability is due to superimposed slowing.

### Waveform shape

Brain rhythms are not pure sinusoidal oscillations, and as such there may be non-linear features of the signal which Fourier analysis fails to capture (Cole and Voytek, 2017). Two metrics previously reported to quantify deviation from sinusoidal shape are rise-decay symmetry and peak-trough symmetry (Cole and Voytek, 2019). Rise-decay asymmetry indicates a sawtooth shape. Peak-trough asymmetry indicates a rounded peak with sharp trough, or *vice versa*. Both of these values are expressed as proportions, with 0.5 indicating equal rise and decay times, or equal peak and trough times. It is unknown whether either of these measures show asymmetry during a spindle. We found that across all cortical sites during NREM, rise-decay was symmetrical, that is equal to 0.5 (LMEM; *β*=0.501, Std. Error= 1e-03,t=0.98, No. channels=357, N=20; Supp. Fig. 1). Spindle cycles were slightly more biased for peaks than troughs (LMEM; *β*=0.509, Std. Error= 2e-03,t=4.4, No. channels=357, N=20; Supp. Fig. 1). It is also unknown whether these measures significantly change during spindles. We found that 16.1% (58/360) of cortical channels showed a significant (paired t-test, p<0.05, FDR adjusted) difference in peak-trough symmetry at the start versus the end of a spindle. Of these 58 significant channels, in 41 peak-trough asymmetry became more peak-biased (p=.0022, binomial test). However, the changes were small, with the average increase in peak-trough asymmetry of these 41 changing from 0.52 to 0.53. The 17 channels that became more trough biased changed on average from 0.49 to 0.48. For rise-decay symmetry, 24.4% (88/360) of channels showed significant differences at the start versus the end. Of these, 38.6% (54/88) on averaged changed from 0.52 to 0.5 and the other 34 cortical sites on average from 0.49 to 0.5. This indicates as the spindle progresses, it becomes slightly more symmetrical in its rise-decay ratio. Overall, these measures are largely symmetrical during a spindle, vary little between cortical regions (Supp. Fig. 1), and less than 25% of sites showed a significant change during a spindle for either measure, however this effect was minor.

### Relationships between spindle characteristics

We also investigated the relationships between spindle frequency and other spindle characteristics. We modeled spindle frequency as a function of spindle duration, intra-spindle variation, frequency change, amplitude, and high gamma power using nested LMEMs, with observations at the spindle level (Table 4). All variables were entered as fixed effects into a full model with nested random effects for channels within patients. Overall, faster spindles were shorter duration (*β*=-0.17, Std. Error= 5.4e-03,t=-30.99, No. spindles=550,475, No. channels=360, N=20), showed lower intra-spindle standard deviation (*β*=-0.43, Std. Error=2.9e-03,t=-147.9), lower amplitude (*β*=-6.7e-03, Std. Error=9.3e-05,t=-71.97), greater high gamma power (*β*=.23, Std. Error=3.7e-03,t=63.3), and more slowing (*β*=-3.9e-03, Std. Error=3.9e-04,t=- 10.2). We found that these results did not change after controlling for differences due to brain region or sleep stage (Supp. Table 1). However, although there were significant associations between frequency and other spindle characteristics, the variance explained by each was low (Table 4). We estimated the variance explained in a separate model for each spindle characteristic using the MuMIn R package (Barton 2020), which provides estimates of the marginal variance explained R^2^_m_, or the variance explained by the fixed effects alone, and the conditional variance explained R^2^_c_, or the variance explained by both fixed and random effects. The R^2^_m_ for duration (0.004), intra-spindle SD (0.036), amplitude (0.027), high gamma power (0.013), and frequency change (6.7e-05) indicate each spindle characteristic only accounts for a small fraction of the variation in frequency. Furthermore, given the average absolute differences of frontal and parietal sites for duration (0.01s), amplitude (9 µV), intra-spindle SD (0.03 Hz), and intra-spindle frequency change (0.46 Hz/s), the predicted absolute frequency differences are 0.002, 0.06, 0.013, and 0.002 Hz, respectively, all far lower than the observed

**Table 4.** Spindle characteristics that covary with overall spindle frequency. Results were unchanged after controlling for sleep stage and regional differences (Supp Table 1). We applied a linear mixed-effects model (LMEM) with nested random effects, channels within patients on 550,475 spindle observations 360 channels, and 20 patients. Estimates, *β*, represent linear slopes of the predictors (rows) on overall spindle frequency with 95% confidence intervals (CI). For example, the Duration *β* of -0.17 indicates that a 1µV increase is associated with a 6.7e-03 Hz decrease in frequency. The columns under ‘Variance Explained’ are the marginal variance explained R^2^_m_, or that explained by the fixed effect alone (e.g. Duration), and the conditional variance explained R^2^_c_, or that explained by both fixed and random effects. The variance explained was estimated for each fixed-effect (row) independently. For example, the additional variance in frequency explained by duration across the entire dataset is very small (0.004) compared to the variance explained by the grouping of spindles within specific channels and patients (0.378).

|  | Frequency (Hz) |  | Variance Explained |  |
| --- | --- | --- | --- | --- |
| | $\beta$ | CI | $R^2_m$ | $R^2_c$ |
| Intercept (Hz) | 12.35 *** | 12.22 – 12.47 |  |  |
| Duration (s) | -0.17 *** | -0.18 – -0.16 | 0.004 | 0.378 |
| Intra-spindle SD (Hz) | -0.43 *** | -0.43 – -0.42 | 0.036 | 0.367 |
| Amplitude ($\mu V$) | -6.7e-03 *** | -6.9e-03 – -6.5e-03 | 0.027 | 0.417 |
| High Gamma Power ($\mu V^2$) | 0.23 *** | 0.23 – 0.24 | 0.013 | 0.370 |
| Frequency Change (Hz/s) | -3.9e-03 *** | -4.7e-03 – -3.1e-03 | 6.7e-05 | 0.370 |

\* $p < 0.05$ \*\* $p < 0.01$ \*\*\* $p < 0.001$

0.64 Hz frontal-parietal frequency difference.

### Spindle co-occurrence

While originally described as a global phenomenon, MEG and intracranial work have identified spindles as primarily local events (Andrillon et al., 2011; Dehghani et al., 2010; Frauscher et al., 2015; Piantoni et al., 2017). We found that indeed the majority of spindles occur at a small proportion of the cortical sites recorded (Figure 3A,B). Typically, individual spindles occurred in only a single or a few channels, with 50% of spindles occurring in under 16% of channels and 75% in under 25% of channels (on average 18 cortical channels per patient were analyzed for spindles). Frontal and parietal sites showed the greatest proportion of multiple channels participating in spindle events (Figure 3A). This effect is also clearly shown in Figure 3C where frontal and parietal sites have the smallest proportion of spindles occurring in only a single channel, and the largest proportion of spindles occurring in four or more channels.

**Figure 3.**
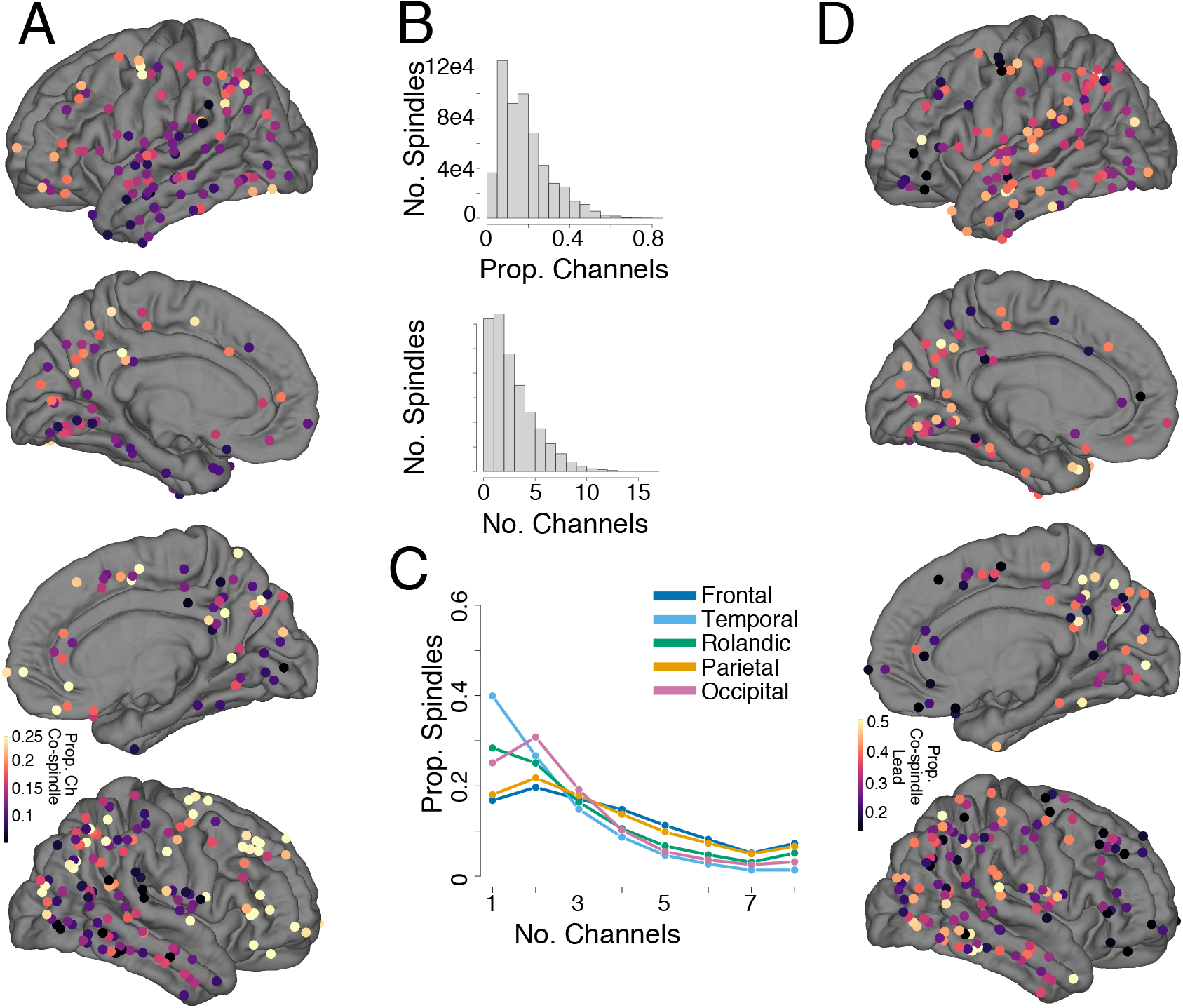
Spindle co-occurrences across the cortex. **A**,**D**. Brain surfaces overlaid with cortical SEEG recording sites. Warmer colors indicate greater values. **A**. The average proportion of channels participating in a spindle at each cortical site; frontal and parietal show the greatest proportion. **B**. Distribution of the proportion of channels participating in a spindle (left) and absolute number of channels participating in a spindle (right) for all spindles from all channels. **C**. For each region, the proportion of spindles that occurred in one to eight channels. **D**. For each channel, the proportion of times a spindle initiated a co-spindling event, i.e. when a spindle co-occurred in at least two channels.

We also found that the proportion of spindles that lead or initiated co-spindling events (e.g. spindles detected in multiple channels) was greatest in parietal and especially medial parietal regions (Figure 3D). In contrast, frontal sites showed the lowest proportion of leading spindles in co-spindle events. During co-spindling events, parietal spindles start earlier than frontal (LMEM; *β* =-81.75 ms, Std. Error = 12.23, t=-6.68, No. spindles=427,662, No. channels= 360, N=20).

Spindles that co-occurred in multiple channels also had unique spindle characteristics (Table 5). Spindles that occurred in successively more channels were faster in frequency (LMEM with nested random effects for channels within patients; No. spindles=554,794, No. channels= 365, N= 20; for single channel vs 6+ channels *β*=0.36 Hz,Std. Error=4e-03,t=86.86), longer duration (for 6+ channels *β*=0.13 s, Std. Error=1e-03,t=126.86), had lower intra-spindle frequency SD (for 6+ channels *β*=-0.14 Hz, Std. Error=2e-03,t=-72.98), and showed significantly greater spindle slowing (for 6+ channels *β*=-0.45 Hz/s, Std. Error=0.01, t=-31.68). These effects remained significant after controlling for sleep stage and regional differences (Supp. Table 2).

**Table 5.** Characteristics of widespread spindles. We applied a linear mixed-effects model (LMEM) with nested random effects, channels within patients on 554,794 spindle observations, 365 cortical channels, and 20 patients. Estimates, *β*, represent contrasts in the dependent variable (columns) between 2 or more channels against a channel. For example, the *β* for 6+ Ch. of 0.36 in the Frequency column indicates that spindles occurring in 6 or more channels have an average frequency 0.36 Hz higher than those that occur in only one channel. Results were unchanged after controlling for sleep stage and regional differences (Supp. Table 2).

| No. Ch | Frequency (Hz) |  | Duration (s) |  | Intra-spindle SD (Hz) |  | Frequency Change (Hz/s) |  |
| --- | --- | --- | --- | --- | --- | --- | --- | --- |
| | $\beta$ | CI | $\beta$ | CI | $\beta$ | CI | $\beta$ | CI |
| 1 | 12.17 *** | 12.04 – 12.29 | 0.66 *** | 0.65 – 0.67 | 0.98 *** | 0.95 – 1.01 | -0.18 ** | -0.30 – -0.06 |
| 2 | 0.09 *** | 0.08 – 0.09 | 0.04 *** | 0.03 – 0.04 | -0.02 *** | -0.03 – -0.02 | -0.11 *** | -0.13 – -0.09 |
| 3 | 0.16 *** | 0.16 – 0.17 | 0.06 *** | 0.06 – 0.06 | -0.05 *** | -0.05 – -0.05 | -0.22 *** | -0.24 – -0.19 |
| 4,5 | 0.25 *** | 0.24 – 0.26 | 0.08 *** | 0.08 – 0.09 | -0.09 *** | -0.09 – -0.09 | -0.36 *** | -0.38 – -0.33 |
| 6+ | 0.36 *** | 0.35 – 0.37 | 0.13 *** | 0.13 – 0.13 | -0.14 *** | -0.14 – -0.14 | -0.45 *** | -0.48 – -0.42 |

\* $p < 0.05$ \*\* $p < 0.01$ \*\*\* $p < 0.001$

### N2 vs N3 differences in spindle characteristics

We compared spindle characteristics between N2 and N3 at the channel level, pooling data from all 20 patients, 357 cortical channels, and 711 unique sleep stage-channel observations. Overall, spindle density was greater in N2 than N3 (LMEM; *β* = 0.39 spindles/min, Std. Error= 0.04,t=9.16). After controlling for regional differences, spindles had slightly longer duration in N2 than N3 (LMEM; *β* = 0.02 s, Std. Error= 2e-03, t=9.18). There were no significant differences in average spindle amplitude between sleep stages (LMEM; *β* = 0.29 µV, Std. Error= 0.21, t=1.35). After controlling for regional differences, N2 estimates of overall frequency were higher than N3 (LMEM; *β* = 0.08 Hz, Std. Error= 0.01, t=5.94) and cortical sites showed a greater inter-spindle standard deviation in frequency for N2 than N3 (LMEM; *β* = 0.05 Hz, Std. Error= 9e-03, t=6.17). While there were no differences between N2 and N3 for intra-spindle SD (LMEM; *β* =5e-03 Hz, Std Error=5e-03, t=-0.97), the frequency range was smaller in N3 (LMEM; *β* =-0.05 Hz, Std Error=0.02, t=-3.32). There were no significant differences in rates of change in frequency between N2 and N3 (LMEM; *β* =-6e-03 Hz/s, Std Error=0.03). Rise-decay symmetry did not differ between N2 and N3 (LMEM; *β* = 4e-04, Std. Error= 7e-04, t=0.56), however peak-trough symmetry was more peak-biased for N3 than N2 (LMEM; *β* = 18e-04, Std. Error= 7e-04, t=2.75). N2 sleep showed a greater amount of spindle co-occurrence than N3 sleep (LMEM; *β* =0.57 channels, Std. Error = 0.01, t=94.72, No. spindles=554,794, No. channels= 365, N=20).

### Left vs right hemisphere differences in spindle characteristics

We compared spindle characteristics between left and right hemispheres at the channel level, pooling data from all 20 patients and 357 cortical channels. During NREM sleep, left and right hemispheres did not differ in: spindle density (LMEM; *β* = -0.14 spindles/min, Std. Error= 0.19,t=-0.73,), spindle amplitude (LMEM; *β* = -1.49 µV, Std. Error= 2.35,t=-0.63), overall frequency (LMEM; *β* = -3e-03 Hz, Std. Error= 0.07,t=-0.05), inter-spindle variation (LMEM; *β* = - 0.02 Hz, Std. Error= 0.02, t=-0.81), within-spindle variation (LMEM; *β* = 0.02 Hz, Std. Error= 0.02,t=1.12), frequency range (LMEM; *β* = 0.05 Hz, Std. Error= 0.05,t=0.97), linear frequency change (LMEM; *β* = 0.04 Hz/s, Std. Error= 0.06, t=0.66), rise-decay symmetry (LMEM; *β* = -6e- 04, Std. Error= 16e-04, t=-0.37), nor in peak-trough symmetry (LMEM; *β* = -4e-03, Std. Error= 2e-03, t=-1.52). We found that spindles occurring in the right hemisphere compared to the left showed greater co-occurrence (LMEM; *β* =0.3 channels, Std. Error = 0.1, t=-3.03, No. spindles= 554,354, No. channels= 363, N=20, and that the average spindle duration was shorter in left versus right hemisphere (LMEM; *β* = -0.01 s, Std. Error= 5.5e-03,t=-2.37).

### Cortical-hippocampal spindle phase locking

In previous work, we found 37% of NC channels showed significant PLV with NC-HC spindles; 67% of these were with posterior hippocampus in N2. The latter channels are shown in Figure 4A, with color indicating the peak PLV during N2 with posterior hippocampal spindles. The PLV among channels with significant HC-NC spindle PLV showed a positive trend with overall spindle frequency after controlling for differences in frequency due to cortical region (LMEM; *β* = 0.78, t=1.92, No. channels= 76, N=12). Intra-spindle frequency SD negatively covaried with PLV after controlling for differences due to cortical region, though this effect was marginal (LMEM; *β* =- 0.2,t= -1.96, No. channels= 76, N=12). Neither spindle duration, frequency change, nor N2 density significantly covaried with PLV after controlling for cortical region (|t|<1.95). However, we previously found 5% of all NC channels showed high NC-HC PLV (peak >0.4), 70% of which were in parietal channels. Post-hoc analyses found after controlling for regional differences, this high PLV subset compared to all other cortical channels were significantly faster (LMEM; *β* =0.57 Hz,t=5.01, No. channels= 341, N=20; Figure 4B), had lower intra-spindle frequency variation (*β* =-0.15 Hz,t=-4.7; Figure 4C), had slightly longer duration (*β* =0.02 s, t=2.82; Figure 4D), and greater spindle density in N2 (*β* =0.76 spindles/min,t=2.48; Figure 4E) as well as in N3 (*β* =0.72 spindles/min,t=2.25). These sites did not differ in linear change in frequency (*β* =-0.1 Hz/s,t=-1.06), inter-spindle frequency standard deviation (*β* =0.03 Hz,t=0.77), or amplitude (*β* =-2.9 µV,t=-0.75). These results did not change when restricting to just parietal channels (LMEM; No. channels=113, N=18). Furthermore, after controlling for the effects of sleep stage, hemisphere, and region, spindles occurring at cortical sites with high PLV also showed greater co-occurrence with spindles across multiple cortical sites (LMEM; *β* = 0.34 channels, Std. Error= 0.13, t=2.5, No. spindles=550,769, No. channels=360, N=20). These analyses demonstrate sites with large HC-NC spindle PLV, predominately parietal sites, have unique spindle characteristics compared to other cortical sites after controlling for regional differences, including faster frequency, lower within-spindle frequency variation, longer duration, greater density, and a greater degree of spindle co-occurrence across cortical sites.

**Figure 4.**
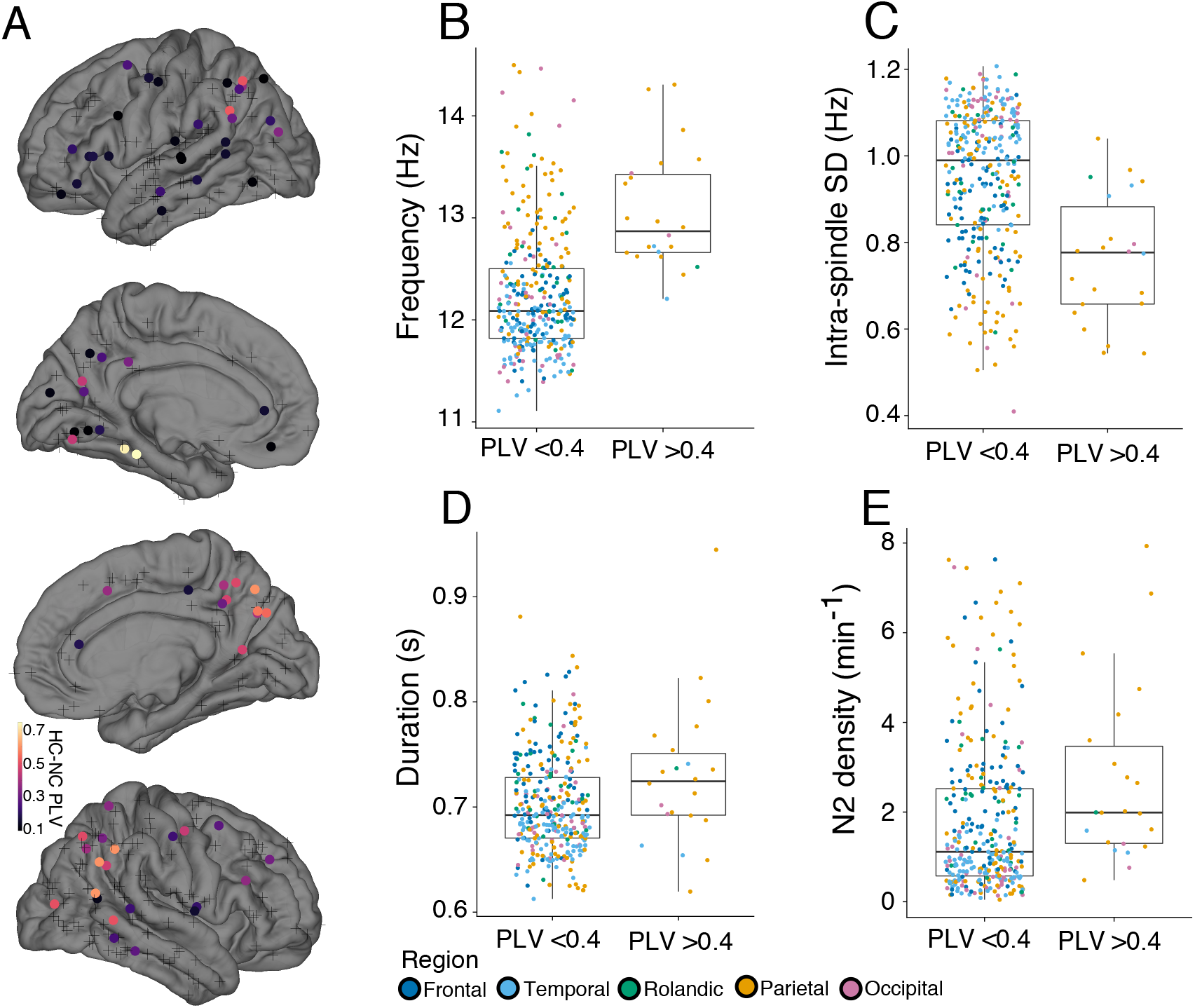
Cortical-hippocampal spindle phase locking. **A**. Channels with significant PLV during spindles with a posterior hippocampal channel during N2, as identified in Jiang et al. 2019b, are indicated as circles, with the peak PLV shown in color. Non-significant channels are displayed as crosses. Areas with the highest PLV are apparent in parietal cortex as well as posterior ventral temporal cortex. We performed post-hoc analyses comparing these high PLV channels (PLV >0.4) and all other cortical channels, including those with non-significant PLV, across several spindle characteristics. **B-E** Each dot is a channel, different colors denote different brain regions. High PLV channels were **B** significantly faster (LMEM; *β* =0.57,t=5.01, No. channels= 341, N=20), **C** had lower intra-spindle frequency variation (*β* =-0.15,t=-4.7), **D** had slightly longer duration (*β* =0.02, t=2.82), and **E** had greater spindle density in N2 (*β* =0.76,t=2.48). These results did not change when restricting to just parietal channels.

### Slower and faster spindles show similar coupling to downstates

Putative evidence for dichotomizing spindles as slow or fast includes that they have different coupling to SOs, wherein frontal, slower spindles precede and faster, centro-parietal spindles follow downstates (Klinzing et al., 2016; Mölle et al., 2011), however recent intracranial work does not support this effect (Gonzalez et al., 2018; Mak-McCully et al., 2017). Studies vary in the cutoff frequency that separates slow from fast spindles, as spindle peak frequencies vary across subjects (Ujma et al., 2015) and some subjects do not show distinct slow and fast spindle peaks in averaged power spectra (Cox et al., 2017; Gennaro and Ferrara, 2003; Mölle et al., 2011; Werth et al., 1997). We chose 12 Hz to demarcate slower and faster spindles, as 12 or 13 Hz is typical (Ayoub et al., 2012; Barakat et al., 2011; Mölle et al., 2011; Schabus et al., 2007), and determined if there is regional variability in whether slow spindles precede downstates and fast spindles follow.

We found that of the 365 channels, 320 had at least 40 spindles starting within +/- 0.5s of cortical downstates. There were significantly more spindles starting 0.5 s after downstates compared with starting before (LMEM; *β* =180.5 spindles, Std. Error= 28.2, t=6.4, No. channels= 320,N=20). Of these 320, 208 showed a significant tendency for spindles to start after (77%, 160 channels) or before (23%, 48 channels) downstates (two-sided binomial test; p<0.05, FDR-adjusted). The probability of spindles starting relative to downstate trough for all regions is shown in Supp. Fig. 2. The majority of significant sites were from frontal (29%,60 channels) and parietal (31%, 64 channels) cortex. All significant channels from frontal and parietal sites, regardless of preferred latency, are shown in Figure 5. To compare differences in coupling of spindles to downstates by frequency, only channels with at least 100 spindles <12 Hz and 100 spindles > 12 Hz within +/- 1s downstates are shown. Gray lines separate patients and each row is a cortical channel. Because the sample sizes of spindles < 12 Hz and > 12 Hz were unbalanced for some channels, we bootstrapped the spindle latency times for both frequency conditions over 10,000 iterations, using the size of the smaller condition. With each iteration, a histogram of spindle start latencies was generated, and the average of these histograms was normalized across bins for each channel (shown as color in Figure 5Ai,Bi). To assess whether there were significant differences between either frequency condition and chance, or between the two conditions, we fit a LMEM on the probability of spindles starting at each time bin and display error bars of standard error (Figure 5 Aii,Bii). The blue dashed line indicates the probability of spindles starting by chance. Time bins where error bars do not include the chance line indicate significance at p<0.05. Both spindles > 12 Hz (pink triangles) and < 12 Hz (gray squares) significantly start following downstate troughs, in both frontal and parietal sites. Whether spindles are overall slower or faster, they show similar temporal relationships to downstates.

**Figure 5.**
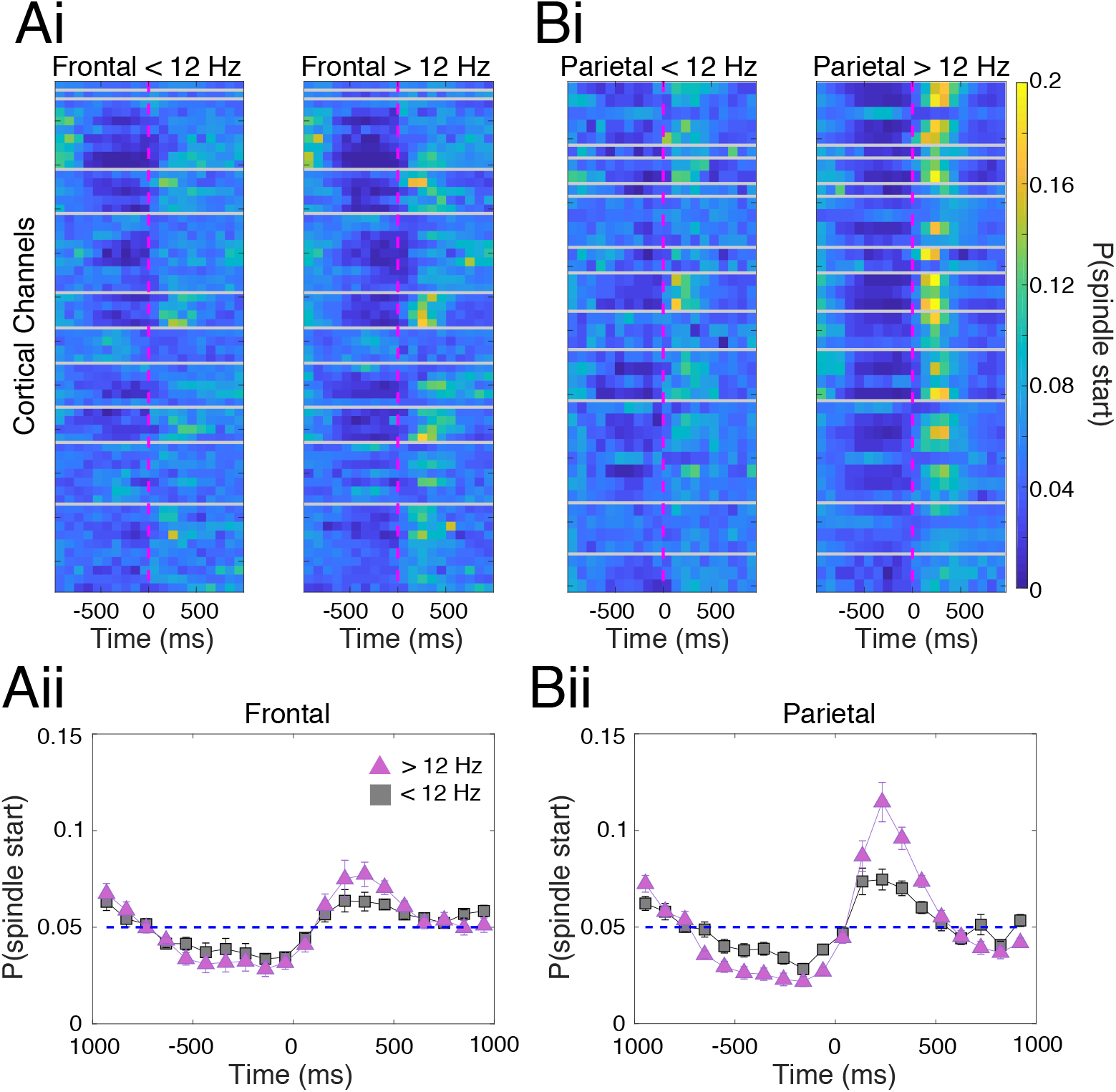
Slower and faster spindles show similar coupling to downstates. **A**,**B** Cortical channels with at least 100 spindles < 12 Hz and 100 spindles > 12 Hz starting within +/-1s of downstate trough are shown. **Ai**,**Bi** Rows indicate cortical channels, gray lines separate patients, and the dashed magenta line indicates time of the downstate trough. Color indicates the probability of spindles starting within that channel. For both regions and frequency groups, there is a greater probability of spindles starting after the downstate trough compared to before. This is statistically assessed in **Aii**,**Bii**, where LMEMs, with patient as random effect, modeled the probability of spindles starting at each time bin for each frequency condition separately, with pink triangles showing spindles >12 Hz and gray squares for spindles < 12 Hz. Error bars reflect standard error, and the blue dashed line indicates the probability of spindles starting by chance. Time bins where error bars do not include the chance line indicate significance at p<0.05. Both frontal and parietal and frequency groups show a decreased probability of starting before, and an increase after downstate trough. Notably, for both **Aii** and **Bii**, there are time bins where spindles > 12 Hz show a greater probability of starting after downstate troughs compared to spindles < 12 Hz.

## Discussion

This study uses sEEG recordings to investigate spindle characteristics including multiple sources of variability in spindle frequency. While spindles were detected in all cortex areas sampled, temporo-occipital sites showed lower density, more variable frequency, and shorter duration spindles. Fronto-parietal regions had the greatest spindle duration, density, and proportion of spindles co-occurring in multiple channels; consistent with them making the largest contribution to EEG. Spindle frequency variability was assessed across spindles within a channel, and across cycles within spindles, then compared to fronto-parietal differences. While, compared to frontal channels, we observed faster overall frequency in Rolandic (0.37Hz) and parietal (0.64 Hz) cortices, the SD across spindles within a site, and across cycles within a spindle, were both larger (0.87 and 0.94 Hz, respectively). We also found that spindles which occurred in multiple channels were faster, showed greater slowing, and had lower intra-spindle variability, thus indicating that local spindles differ from more global events. Previous work identified a subset of parietal channels with high phase locking to posterior hippocampal spindles (Jiang et al., 2019b); here we describe how these cortical sites have spindles with faster frequency, longer duration, lower intra-spindle variability, and greater density. Although our findings are from epileptic patients and not a healthy population, we analyzed a large number of patients (n=20) with unique epileptiform etiologies, had broad cortical coverage, excluded sleep periods or channels with pervasive epileptic activity, and analyzed spindles whose appearance as well as other spindle characteristics such as density were within healthy ranges.

The dichotomy of parietal-fast and frontal-slow spindle types was introduced based on EEG (Gibbs and Gibbs, 1950). However, most intracranial work reports a gradient of spindle frequencies (Frauscher et al., 2015; Peter-Derex et al., 2012; Piantoni et al., 2017), with an exception being Andrillon et al. 2011. Our findings suggest a modification to the model of spindles as dichotomous slow and fast systems. We found individual locations exhibit both faster and slower spindles, and that there is a large overlap in the distribution of frequencies between frontal and parietal sites (Figure 1A). The width of frequency distributions at individual sites was broader in Rolandic-parietal-occipital than frontal-temporal sites (Figure 2A), as previously reported along medial structures (Andrillon et al., 2011).

In addition to the variability in spindle frequency at individual sites, there is also substantial variability within each spindle. We found the fastest and slowest cycles within a spindle on average differed by 2.7 Hz (Table 3, Figure 2C), greater in temporal-occipital sites, and lower at frontal-parietal (Figure 2B,2C). This variability includes a systematic decrease in cycle frequency over the course of spindles, previously observed with EEG (O’ Reilly and Nielsen, 2014; Souza et al., 2016) and MEG (Dehghani et al., 2011a). In our study, we found that 46% of recordings from medial and lateral cortex had a significant change in linear frequency; of these, nearly all (95%) slowed, with an overall average of -0.34 Hz/s, and greatest in frontal cortex at -0.74 Hz/s. In sum, these different sources of variability (inter-spindle, intra-spindle expressed as frequency range, and intra-spindle slowing) all vary by cortical region, and in the majority of channels, exceed the net 0.64 Hz frontal-parietal difference (Figure 2, see black triangles on color bars).

MEG and intracranial recordings have shown that spindles are largely local phenomena (Andrillon et al., 2011; Dehghani et al., 2010; Frauscher et al., 2015; Piantoni et al., 2017). Here, we found most spindles occurred in only a single or a few channels (Figure 3B). Co-occurring spindles apparently underly scalp EEG. Specifically, intracranial work found that scalp spindles were associated with asynchronous sigma activity, predominantly at frontoparietal sites (Frauscher et al., 2015), and simultaneous MEG/EEG found spindles detected in both modalities had a 66% increase in the number of MEG sensors involved in the spindle, especially over frontal sensors (Dehghani et al., 2011b). Similarly, we found that frontal and parietal sites showed the greatest proportion of spindles occurring in multiple channels (Figure 3A,C). We also replicated the phenomenon, previously observed at the scalp (Dehghani et al., 2011a; Mölle et al., 2011) and intracranially (Andrillon et al., 2011), that spindles at central-posterior sites precede anterior spindles (Figure 3D). Furthermore, we found that spindles occurring across multiple channels exhibit faster overall frequency, lower within-spindle frequency variability, and much greater spindle slowing (Table 5). These differences could reflect stereotyped, global propagation patterns, such as rotating waves (Muller et al., 2016). Our finding that parietal spindles precede frontal spindles agrees with the previously observed temporal-parietal-frontal progression of spindle cycles. However these patterns were identified with electrocorticography and our bipolar transcortical sEEG recordings are unambiguously local but more irregularly spaced across the cortex.

Communication between hippocampal and cortical rhythms could serve as a substrate for restructuring recent experiences into long-term memory. Previous work has shown that ripples in the hippocampus are phase locked to hippocampal spindles (Jiang et al., 2019b; Staresina et al., 2015), which in turn are phase-locked with inferior parietal spindles (Jiang et al., 2019b), in locations that are activated in recollective experiences (Gilmore et al., 2015; Hoppstädter et al., 2015). We found that these cortical sites also had unique spindle characteristics, including faster frequency, lower intra-spindle frequency variation, longer duration, and greater density (Figure 4). We speculate that these spindles could play a unique role in processing detailed, episodic information. Considering spindles at these sites spread more widely, they could coordinate or drive spindle dynamics at other cortical structures.

The assertion that slow spindles precede downstates and fast spindles follow is often cited as evidence for their distinct generating mechanisms (Klinzing et al., 2016; Mölle et al., 2011; Timofeev and Chauvette, 2013). Replicating our previous work (Gonzalez et al., 2018; Mak-McCully et al., 2017), we found that spindles were significantly more likely to start after downstates across all cortical sites. We showed that within the same channels, and in both frontal and parietal sites, spindles above and below 12 Hz both follow cortical downstates (Figure 5). Faster spindles showed a greater likelihood of initiating after downstates than slower spindles, especially in parietal cortex (Figure 5Bii). Overall, these findings affirm spindles, regardless of frequency, show similar temporal relationships to downstates and are consistent with slower and faster spindles existing along a continuum instead of arising from distinct neurophysiological generators.

The current study found that spindles vary considerably across multiple characteristics, including duration, amplitude, density, spread, onset, and relation to other sleep waves and structures, as well as in the variability of these characteristics. This variation was observed between individual cycles of spindles, between different spindles recorded from the same electrode derivation, between different electrodes in the same cortical area, and of course between different cortical areas. However, the variation between cortical areas was usually smaller than other sources of variation. Although there was sometimes association between different characteristics, this was not systematic or dichotomous. Thus, in contrast to the conception of spindles as comprising two distinct categories differing in location, frequency, relation to downstates, and presumably all other characteristics, we observe a broad range of characteristics in each structure, electrode, and spindle, with variable relationships. Our data indicates that to the extent that regular differences can be discerned in anterior versus posterior EEG recordings, they reflect the sum of many variable cortical generators rather than two monolithic and uniform generators.

This reconceptualization of spindles as varied and heterogenous has implications for interpreting the finding that fast but not slow spindles are involved in memory consolidation (Barakat et al., 2011; Mölle et al., 2011). However, these studies conflated spindle frequency with sensors (i.e. did not consider faster frontal spindles or slower central-parietal spindles), and dichotomized frequency as either slow or fast. Analyzing memory consolidation as a function of frequency treated as a continuous variable in both frontal and posterior spindles would be needed to examine if spindle frequency *per se* moderates learning and memory. Here, we found faster spindles are associated with greater high gamma (Table 4) and are more tightly associated with initiating on the down-to-upstate transition (Figure 5). Since spindles occurring on down-to-upstate transitions are associated with greater calcium influx to layer 2/3 mouse pyramidal neurons, compared to spindles alone (Niethard et al., 2018), faster spindles could be better suited for facilitating cortical plasticity. Considering spindles as heterogenous also has implications for using spindles as biomarkers in psychiatric disorders such as schizophrenia (Ferrarelli et al., 2007; Manoach and Stickgold, 2019). For example, deficits in faster spindles would not imply an exclusive targeting of central-parietal sites for observation or treatment.

In summary, we found that both frontal and parietal cortex show large numbers of both ‘fast’ and ‘slow’ spindle cycles, and both types occur mainly following downstates. Other characteristics vary quasi-independently, clearly indicating that cortical spindle generators form a continuum. This reconceptualization may inform future efforts to relate spindles to memory consolidation and clinical disorders.

## Acknowledgements

We thank Burke Rosen, Charlie Dickey, Adam Niese, Jacob Garrett, Maxim Bazhenov, Terrence Sejnowski for feedback and support. We also thank Scott Cole and Tammy Tran for providing assistance and custom code for waveform shape analysis.

