## Supplemental material for "Human spindle variability"

**Supplementary Material**

|  | **Frequency** | |
| --- | --- | --- |
| *Predictors* | *Estimates* | *CI* |
| Intercept (Hz) | 12.07 ^***^ | 11.91 – 12.24 |
| Duration (s) | -0.19 ^***^ | -0.20 – -0.18 |
| Intra-spindle SD (Hz) | -0.43 ^***^ | -0.43 – -0.42 |
| Amplitude (µV) | -0.01 ^***^ | -0.01 – -0.01 |
| High Gamma Power (uV^2^) | 0.24 ^***^ | 0.23 – 0.25 |
| Frequency Change (Hz/s) | -3.7e-03 ^***^ | -4.4e-03 – -2.9e-03 |
| N3 | 0.11 ^***^ | 0.11 – 0.12 |
| Temporal | -0.08 | -0.23 – 0.07 |
| Rolandic | 0.44 ^***^ | 0.25 – 0.64 |
| Parietal | 0.72 ^***^ | 0.56 – 0.88 |
| Occipital | 0.29 ^**^ | 0.09 – 0.49 |
| ** p<0.05   ** p<0.01   *** p<0.001* | | |

Supp. Table 1. Spindle characteristics that covary with overall spindle frequency. Sleep stage and brain region have been included as covariates. Here, frontal spindles during N2 serve as the reference, or ‘intercept’. We applied a LMEM with nested random effects, channels within patients on 550,475 spindle observations 360 channels, and 20 patients. Estimates, $\beta$ ,represent linear slopes of the predictors (rows) on overall spindle frequency with 95% confidence intervals (CI). Results remain unchanged compared to Table 4.

|  | **Frequency (Hz)** | | **Duration (s)** | | **Intra-spindle SD**  **(Hz)** | | | | **Frequency Change**  **(Hz/s)** |
| --- | --- | --- | --- | --- | --- | --- | --- | --- | --- |
| *Predictors* | $\beta$ | *CI* | $\beta$ | *CI* | $\beta$ | *CI* | $\beta$ | *CI* | |
| Intercept  (Hz) | 11.91 ^***^ | 11.75 – 12.08 | 0.66 ^***^ | 0.65 – 0.67 | 0.95 ^***^ | 0.91 – 0.99 | -0.56 ^***^ | -0.70 – -0.42 | |
| N3 | -0.07 ^***^ | -0.08 – -0.07 | -0.03 ^***^ | -0.03 – -0.03 | -0.01 ^***^ | -0.01 – -0.01 | 0.06 ^***^ | 0.04 – 0.07 | |
| Temporal | -0.08 | -0.24 – 0.08 | -0.02 ^**^ | -0.03 – -0.01 | 0.11 ^***^ | 0.07 – 0.16 | 0.58 ^***^ | 0.45 – 0.71 | |
| Rolandic | 0.45 ^***^ | 0.24 – 0.65 | 0.01 | -0.01 – 0.02 | 0.03 | -0.02 – 0.09 | 0.30 ^***^ | 0.14 – 0.47 | |
| Parietal | 0.74 ^***^ | 0.57 – 0.91 | 9.5e-04 | -0.01 – 0.01 | -0.06 ^**^ | -0.11 – -0.02 | 0.46 ^***^ | 0.32 – 0.59 | |
| Occipital | 0.27 ^*^ | 0.06 – 0.47 | -4e-03 | -0.02 – 0.01 | 0.08 ^**^ | 0.02 – 0.13 | 0.55 ^***^ | 0.38 – 0.72 | |
| 2 Ch | 0.08 ^***^ | 0.08 – 0.09 | 0.03 ^***^ | 0.03 – 0.04 | -0.02 ^***^ | -0.03 – -0.02 | -0.11 ^***^ | -0.13 – -0.09 | |
| 3 Ch | 0.16 ^***^ | 0.15 – 0.17 | 0.06 ^***^ | 0.06 – 0.06 | -0.05 ^***^ | -0.05 – -0.05 | -0.21 ^***^ | -0.24 – -0.19 | |
| 4,5 Ch | 0.24 ^***^ | 0.23 – 0.25 | 0.08 ^***^ | 0.08 – 0.08 | -0.09 ^***^ | -0.09 – -0.09 | -0.35 ^***^ | -0.37 – -0.33 | |
| 6+ Ch | 0.34 ^***^ | 0.34 – 0.35 | 0.12 ^***^ | 0.12 – 0.12 | -0.14 ^***^ | -0.14 – -0.14 | -0.44 ^***^ | -0.47 – -0.41 | |
| ** p<0.05   ** p<0.01   *** p<0.001* | | | | | | | | | |

Supp. Table 2. We applied a LMEM with nested random effects, channels within patients on 550,769 spindles, 360 cortical channels, and 20 patients. Estimates, $\beta$ ,represent contrasts in the dependent variable (columns) between 2 or more channels against the reference or ‘Intercept’, which are spindles occurring in one channel in frontal cortex during N2. Results remain unchanged compared to Table 5.

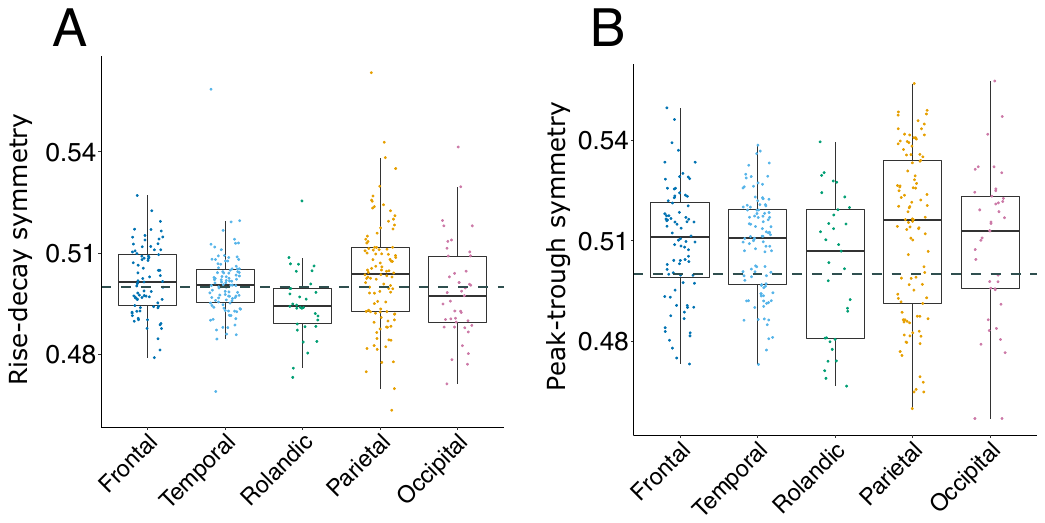

Supp. Fig. 1. Waveform shape across regions. **A** Average rise-decay symmetry for each bipolar recording (indicated as dots), divided by cortical region. **B** Same as A, for peak-trough symmetry. Dashed lines indicate 0.5, or symmetrical measures in rise-decay and peak-trough.

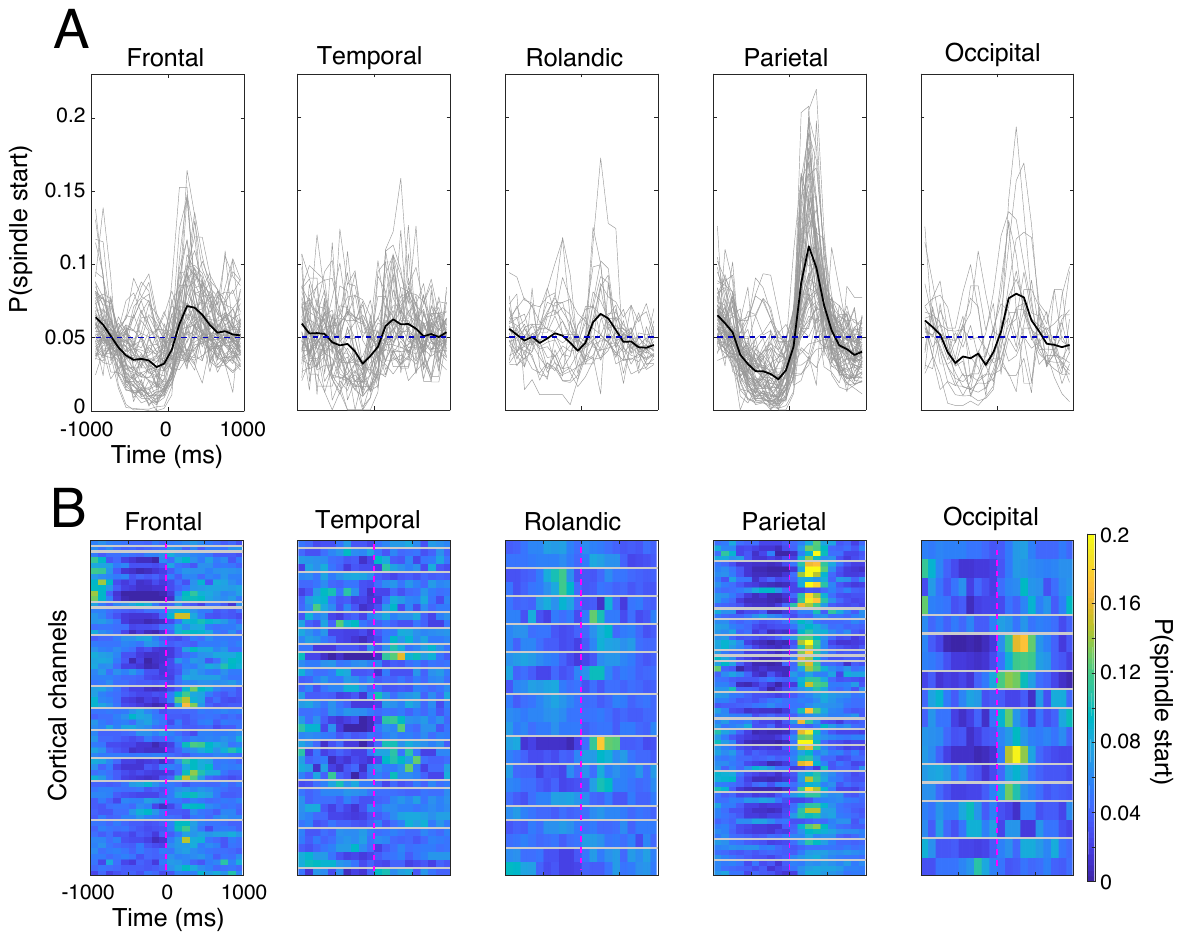

Supp. Fig. 2. Times of spindles relative to downstate troughs. **A** Each gray line corresponds to a bipolar, cortical recording, with black lines indicating average across channels. **B.** Each row is a cortical channel, with gray lines separating different patients and magenta lines indicating the time of downstate trough. The y-axis in **A** and color in **B** indicate the probability of spindles starting in a particular time bin. For each region, all spindles starting within +/- 1s of the downstate trough are included. Averages for most regions show an increase in probability after the downstate trough. The most consistent effect across channels is that spindles have the highest probability of starting after the downstate trough, most apparent in frontal and parietal cortex.
